# Deep learning detection of informative features in tau PET for Alzheimer’s disease classification

**DOI:** 10.1101/2020.07.20.212852

**Authors:** Taeho Jo, Kwangsik Nho, Shannon L. Risacher, Andrew J. Saykin, for the Alzheimer’s Neuroimaging Initiative

**Affiliations:** Department of Radiology and Imaging Sciences, Center for Neuroimaging, Indiana University School of Medicine, Indianapolis, IN, United States; Indiana Alzheimer’s Disease Research Center, Indiana University School of Medicine, Indianapolis, IN, United States; Indiana University Network Science Institute, Bloomington, IN, United States

**Author notes:** Data used in preparation of this article were obtained from the Alzheimer’s Disease Neuroimaging Initiative (ADNI) database (adni.loni.usc.edu). As such, the investigators within the ADNI contributed to the design and implementation of ADNI and/or provided data but did not participate in analysis or writing of this report. A complete listing of ADNI investigators can be found at: http://adni.loni.usc.edu/wp-content/uploads/how_to_apply/ADNI_Acknowledgement_List.pdf. Email addresses: TJ, KN, SLR, AJS.

## Abstract

**Background:** Alzheimer’s disease (AD) is the most common type of dementia, typically characterized by memory loss followed by progressive cognitive decline and functional impairment. Many clinical trials of potential therapies for AD have failed, and there is currently no approved disease-modifying treatment. Biomarkers for early detection and mechanistic understanding of disease course are critical for drug development and clinical trials. Amyloid has been the focus of most biomarker research. Here, we developed a deep learning-based framework to identify informative features for AD classification using tau positron emission tomography (PET) scans.

**Methods:** We analysed [^18^F]flortaucipir PET image data from the Alzheimer’s Disease Neuroimaging Initiative (ADNI) cohort. We first developed an image classifier to distinguish AD from cognitively normal (CN) older adults by training a 3D convolutional neural network (CNN)-based deep learning model on tau PET images (N=132; 66 CN and 66 AD), then applied the classifier to images from individuals with mild cognitive impairment (MCI; N=168). In addition, we applied a layer-wise relevance propagation (LRP)-based model to identify informative features and to visualize classification results. We compared these results with those from whole brain voxel-wise between-group analysis using conventional Statistical Parametric Mapping (SPM12).

**Results:** The 3D CNN-based classification model of AD from CN yielded an average accuracy of 90.8% based on five-fold cross-validation. The LRP model identified the brain regions in tau PET images that contributed most to the AD classification from CN. The top identified regions included the hippocampus, parahippocampus, thalamus, and fusiform. The LRP results were consistent with those from the voxel-wise analysis in SPM12, showing significant focal AD associated regional tau deposition in the bilateral temporal lobes including the entorhinal cortex. The AD probability scores calculated by the classifier were correlated with brain tau deposition in the medial temporal lobe in MCI participants (r=0.43 for early MCI and r=0.49 for late MCI).

**Conclusion:** A deep learning framework combining 3D CNN and LRP algorithms can be used with tau PET images to identify informative features for AD classification and may have application for early detection during prodromal stages of AD.

## Introduction

The accumulation of hyperphosphorylated and pathologically misfolded tau protein is one of the cardinal and most common features in Alzheimer’s disease (AD) [1-5]. The amount and spatial distribution of abnormal tau, seen pathologically as neurofibrillary tangles in brain, is closely related to the onset of cognitive decline and the progression of AD. The identification of morphological phenotypes of tau on *in vivo* neuroimaging may help to differentiate mild cognitive impairment (MCI) and AD from cognitively normal older adults (CN) and provide insights regarding disease mechanisms and patterns of progression [6-9].

Deep learning has been used in a variety of applications in response to the increasingly complex and growing amount of medical imaging data [10-12]. Significant efforts have been made regarding the application of deep learning to AD research, but predicting AD progression through deep learning using neuroimaging data has focused primarily on magnetic resonance imaging (MRI) and/or amyloid positron emission tomography (PET) [10, 13]. However, MRI scans cannot visualize molecular pathological hallmarks of AD, and amyloid PET cannot, without difficulty, visualize the progression of AD due to the accumulation of amyloid-β early in the disease course with a plateau in later stages [14, 15].

The presence and location of pathological tau deposition in the human brain is well established [2, 3, 5]. Braak and Braak [5] analyzed AD-related neuropathology and generated a staging algorithm to describe the tau anatomical distribution [6, 8, 16, 17]. Their results have been confirmed by subsequent studies showing that the topography of tau corresponds with the pathological stages of neurofibrillary tangle deposition. Cross-sectional autopsy data shows that AD-related tau pathology may begin with tau deposition in the medial temporal lobe (Braak stages I / II), then moves to the lateral temporal cortex and part of the medial parietal lobe (stage III / IV), and eventually to broader neocortical regions (V / VI).

In this study, we developed a novel deep learning-based framework that identifies the morphological phenotypes of tau deposition in tau PET images for the classification of AD from CN. Application of CNN to tau PET is novel as the spatial characteristics and interpretation are quite different compared to amyloid PET, FDG PET or MRI. In particular, the regional location and topography of tau PET signal is considered to be more important than for other molecular imaging modalities. This has implications for how CNN interacts with the complex inputs as well as for visualization of informative features. The deep learning-derived AD probability scores were then applied to prodromal stages of disease including early and late mild cognitive impairment (MCI).

## Materials and Methods

### Study participants

All individuals included in the analysis were participants in the Alzheimer’s Disease Neuroimaging Initiative (ADNI) cohort [18, 19]. A total of 300 ADNI participants (N=300; 66 CN, 66 AD, 97 early mild cognitive impairment (EMCI), and 71 late MCI (LMCI)) with [^18^F]flortaucipir PET scans were available for analysis [1]. Genotyping data were also available for all participants [19]. Informed consent was obtained for all subjects, and the study was approved by the relevant institutional review board at each data acquisition site.

### Alzheimer’s Disease Neuroimaging Initiative (ADNI)

ADNI is a multi-site longitudinal study investigating early detection of AD and tracking disease progression using biomarkers (MRI, PET, other biological markers, and clinical and neuropsychological assessment) [1]. Demographic information, PET and MRI scan data, and clinical information are publicly available from the ADNI data repository (http://www.loni.usc.edu/ADNI/).

### Imaging processing

Pre-processed [^18^F]flortaucipir PET scans (N=300) were downloaded from the ADNI data repository, one scan per individual. Scans were normalized to Montreal Neurologic Institute (MNI) space using parameters generated from segmentation of the T1-weighted MRI scan in Statistical Parametric Mapping v12 (SPM12) (www.fil.ion.ucl.ac.uk/spm/). Standard uptake value ratio (SUVR) images were then created by intensity-normalization using a cerebellar crus reference region.

### Deep learning method for AD classification

Deep learning is a subset of machine learning that has been applied in various fields [20, 21]. Deep learning uses a back-propagation procedure [22], which utilizes gradient descent for the efficient error functions and gradient computing [10, 23-26]. The weights are updated after the initial error value is calculated by the least squares method until the differential value becomes 0, as in the following formula:

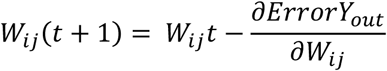

Here, *W*_*ij*_*t* is a current weight of neuron *j* in layer *i*, and *W*_*ij*_(*t* + 1) is the next. *ErrorY*_*out*_ is the sum of errors that are known through the given data. *W*_*ij*_ can be calculated by the chain rule as follows:

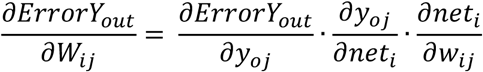

*Net* is a sum of weights and bias, and *Y*_*oj*_ is an output of neuron *j*. Convolutional Neural Network (CNN) is a method of inserting convolution and pooling layers to the basic structure of this neural network to reduce complexity. Since CNN is widely used in the field of visual recognition, we used a CNN method for the classification of AD from CN [27]. The overall architecture of 3D CNN that we used is shown in Fig. 1. To avoid excessive epochs that can lead to overfitting, an early stopping method was applied to cease training if the model did not show improvement over 10 iterations. The learning rate of 0.0001 and Adam, a first-order gradient based probabilistic optimization algorithm [28] with a batch size of 4, were used for training a model. Feature maps (8, 16, and 32 features) were extracted from three hidden layers, with Maxpool3D and BatchNorm3D applied to each layer [29]. Dropout (0.4) was applied to the second and third layers. The training model was applied to the validation set, and each image was determined as AD or CN. Five-fold cross validation was applied, where 60% of the entire data set was used for training, 20% for testing, and 20% for validating. Training images were augmented by three criteria: flipping the image data, shifting the position within two voxels, and shifting the position simultaneously with the flip. Each fold was repeated four times for a robustness check, and the mean accuracy of the four repeats was used as the final accuracy. Pytorch 1.0.1 was used to design neural networks and load pre-trained weights, and all of the programs were run on Python 3.5.

**Fig. 1.**
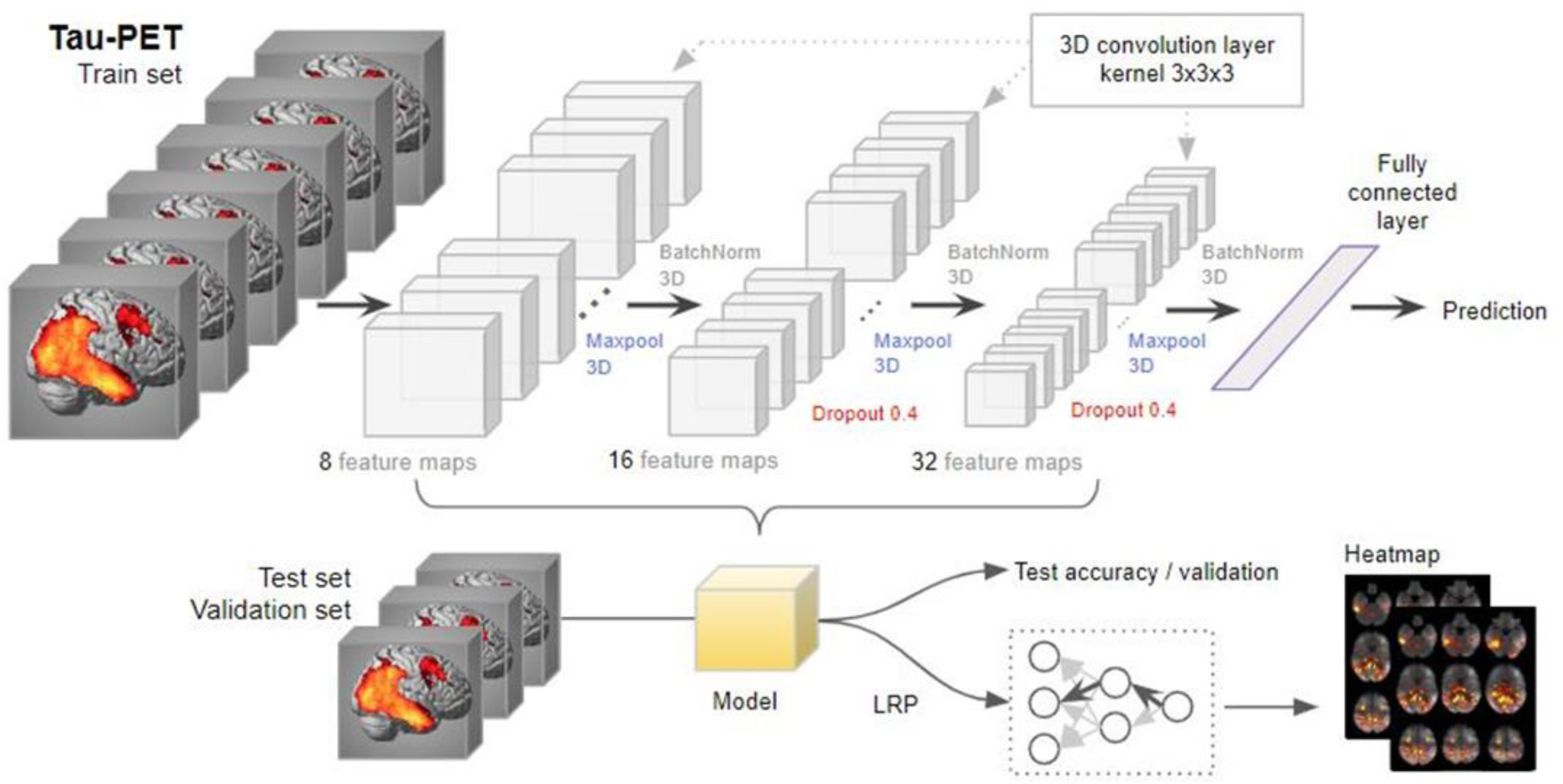
3D convolutional neural network (3D-CNN)-based and layer-wise relevance propagation (LRP)-based framework for the classification of Alzheimer’s disease and the identification of informative features.

### Application of the AD-CN derived classification model to MCI

After an AD classification model was constructed using AD and CN groups, the model was applied to the tau PET scans from the MCI participants to calculate AD probability scores. The AD probability scores were distributed from 0 to 1, and individuals with AD probability scores closer to 1 were classified as having AD characteristics, and individuals with scores closer to 0 were classified as having CN characteristics.

### Identification of informative features for AD classification

We applied a layer-wise relevance propagation (LRP) algorithm to identify informative features and visualize the classification results [30, 31]. The LRP algorithm is used to determine the contribution of a single pixel of an input image to the specific prediction in the image classification task (for full details of the LRP algorithm, see [30]).

The output *x*_*j*_ of a neuron *j* is calculated by a nonlinear activation function *g* and function *h* such as

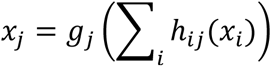

If the relevance score R of the *j* neuron in the layer *l* + 1 sets to 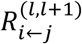, the relevance score R sent to the neuron *i* in the layer *l* will be represented as 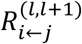. So, the input value of the neuron *j* can be expressed as the following equation:

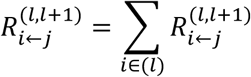

Bach, et al. [30] proposed the following formula for calculating 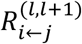:

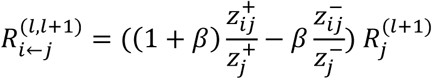

Here, 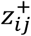 represents the positive input that the node *i* contributes to the node *j*, and 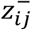 represents the negative input. The variable β that ranges from 0 to 1 controls the inhibition of the relevance redistribution. A larger β value (e.g. β = 1) makes the heat map clearer [31]. In this experiment, we set β=1.

### Whole-brain imaging analysis

A voxel-wise whole brain analysis to identify brain regions in the tau PET SUVR images showing significantly higher tau deposition in AD relative to CN was conducted in SPM12. The analysis was masked for grey plus white matter. The voxel-wise family-wise error (FWE) correction was applied at p < 0.05, with a cluster size of ≥ 50 voxels for adjustment for multiple comparisons.

## Results

In the analysis, 300 ADNI participants (66 CN, 66 AD, 97 EMCI, and 71 LMCI) who had baseline tau PET scans were used. Sample demographics were given in **Table 1**.

**Table 1.** Demographic information

|  | AD | CN | EMCI | LMCI | Total |
| --- | --- | --- | --- | --- | --- |
| **n** | 66 | 66 | 97 | 71 | 300 |
| **Age** | 76.6 | 69.3 | 73.4 | 73.4 | 73.2 |
| **(SD)** | (8.9) | (5.4) | (7.5) | (8.0) | (7.5) |
| **% male** | 56.1% | 40.0% | 37.9% | 66.7% | 50.2% |
| **Education** | 15.8 | 17.2 | 16.3 | 16.4 | 16.4 |
| **(SD)** | (2.5) | (2.1) | (2.8) | (2.5) | (2.5) |
| **% amyloid+** | 90.9% | 27.3% | 40.9% | 48.5% | 51.9% |
| **% ApoE4 carriers** | 51.5% | 25.7% | 35.1% | 28.2% | 35.1% |

### Classification of AD from CN

We developed an image classifier to distinguish AD from CN by training a 3D CNN-based deep learning model on tau PET images. As the number of individuals with AD who had tau PET data were smaller than those of CN, we chose the same number of CN randomly (66 CN) in order to train a classifier with a balanced dataset. In the binary classification problem, it is a well-known issue that detecting disease when the majority of the applicants are healthy, the majority group may be referred as cases, causing biased classification[32]. So we used a random under-sampling (RUS) method to decrease samples from the majority group. All analyses were performed using five-fold cross-validation to reduce the likelihood of overfitting. Ultimately, cross validation in a novel independent data set will be important when such data becomes available. The classification accuracy is shown in **Table 2**. Our deep learning-based classification model of AD from CN yielded an average accuracy of 90.8% and a standard deviation of 2% from five-fold cross-validation (**Table 2**).

**Table 2.** Results of 5-fold cross validation. The numbers in parentheses are the training images after applying augmentation. In addition to the test set, a separate set of 20% independent validations were generated for each fold to ensure the robustness of the experiments. The experiment was repeated four times for each fold (Acc. r1 ∼ r4), and the mean accuracy was considered as the final accuracy of the folds. If the accuracy improvement for the test set did not occur up to 10 times, the training was stopped (epoch).

|  | Train | Test | Val | Acc. r1 | epoch | Acc. r2 | epoch | Acc. r3 | epoch | Acc. r4 | epoch | Mean acc. | SD |
| --- | --- | --- | --- | --- | --- | --- | --- | --- | --- | --- | --- | --- | --- |
| fold1 | 78 (312) | 28 | 26 | 85.7 | 23 | 92.9 | 24 | 89.3 | 21 | 85.7 | 27 | 88.4 | 3.0 |
| fold2 | 78 (312) | 28 | 26 | 96.2 | 20 | 96.2 | 21 | 92.3 | 21 | 88.5 | 22 | 93.3 | 3.2 |
| fold3 | 80 (320) | 26 | 26 | 100 | 35 | 92.3 | 28 | 88.5 | 27 | 88.5 | 30 | 92.3 | 4.7 |
| fold4 | 80 (320) | 26 | 26 | 92.3 | 24 | 88.5 | 31 | 88.5 | 38 | 88.5 | 29 | 89.4 | 1.7 |
| fold5 | 80 (320) | 26 | 26 | 92.3 | 50 | 80.8 | 36 | 96.2 | 35 | 92.3 | 34 | 90.4 | 5.8 |
| AVG: |  |  |  |  |  |  |  |  |  |  |  | 90.8 | 2.0 |

### Identification of informative features for AD classification

The LRP algorithm generated relevance heatmaps in the tau PET image to identify which brain regions play a significant role in a deep learning-based AD classification model. After selecting an AD classification model with the highest accuracy in each fold, we generated five heatmaps and selected the top ten regions with the highest contribution. **Figure 2A** shows a visualization of the relevance heatmap in three orientations of our 3D CNN-based classification of AD from CN. The heatmap displays the primary brain regions that contributed to the classification, color-coded with increasing values from red to yellow. The colored regions in the heatmap include the hippocampus, parahippocampal gyrus, thalamus, and fusiform gyrus (**Fig. 2A**). For comparison with our 3D CNN-based LRP results, **Figure 2B** shows the results of whole brain voxel-wise analysis in SPM12 to identify brain regions where there are significant differences between AD and CN in brain tau deposition (FWE corrected p-value < 0.05; minimum cluster size (k) = 50). AD had significantly higher tau deposition in widespread regions including the bilateral temporal lobes with global maximum differences in the right and left parahippocampal regions, compared to CN (**Fig. 2B**). The informative regions for AD classification in the LRP results is very similar to those found using SPM12, but the 3D CNN-based LRP identified smaller focal regions.

**Fig 2.**
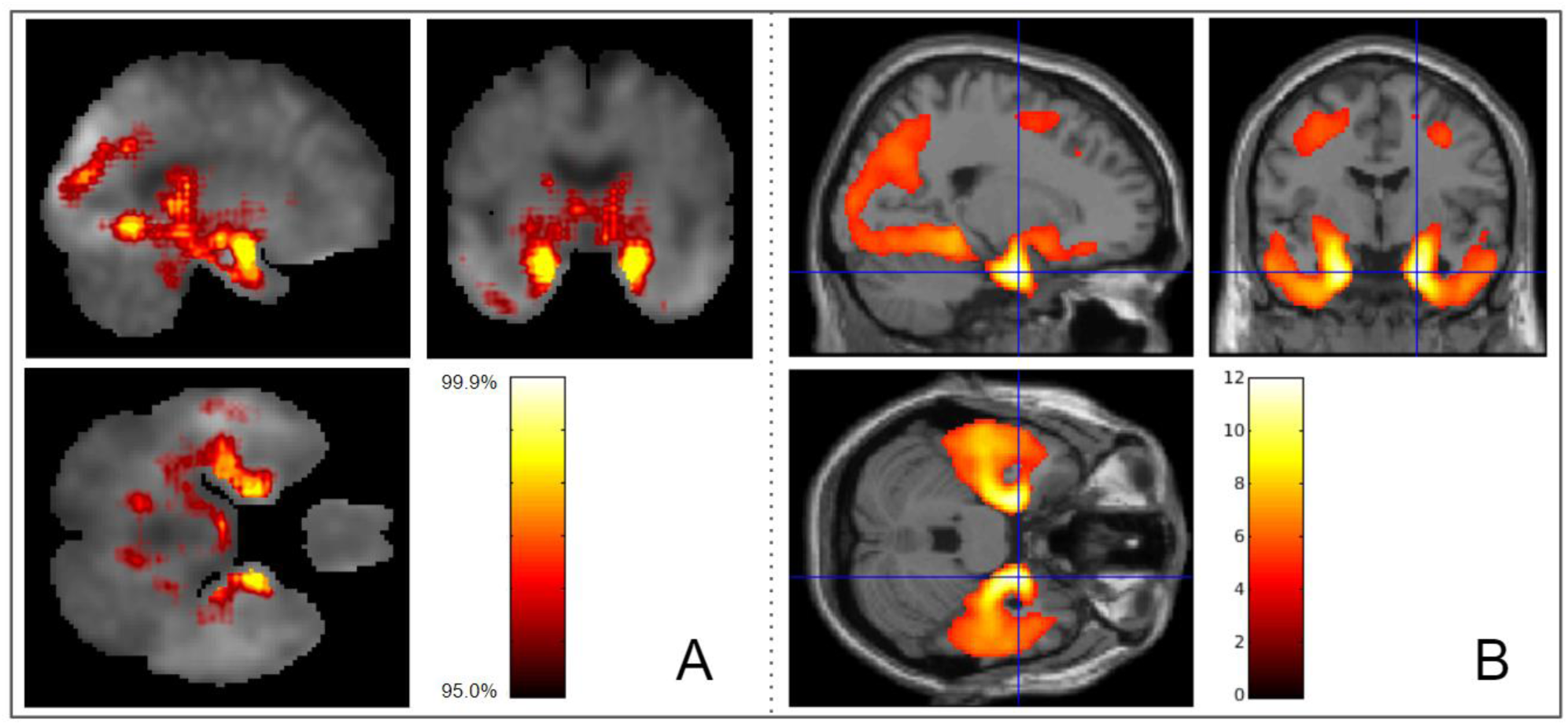
Heatmaps of 3D-CNN classifications compared to voxel-wise group difference maps between AD and CN participant groups. **A**. Relevance heatmaps of 3D-CNN classification of AD and CN. The bright areas represent the regions that most contribute to the CN/AD classification in CNN. Selected regions with the highest contribution include the hippocampus, parahippocampal gyrus, thalamus, fusiform gyrus, and diencephalon. **B**. SPM maps show similar regions of the brain as the 3D-CNN maps where tau deposition is significantly higher in the AD group compared to the CN group (Voxel-wise FWE-corrected p-value < 0.05; minimum cluster size (k) = 50).

### Classification of MCI based on the AD-CN classification model

We calculated the AD probability scores of MCI participants (97 EMCI and 71 LMCI, separately) using the classification model generated above. **Figure 3A** and **Figure 3B** show scatter plots between the AD probability scores of EMCI and LMCI, respectively, with bilateral mean tau deposition in the medial temporal lobe (includes the entorhinal cortex, fusiform, and parahippocampal gyri). The correlation coefficients were R = 0.43 for EMCI and R = 0.49 for LMCI, with greater tau deposition levels in the medial temporal lobe associated with higher AD probability scores. Figure 3C and Figure 3D show mean tau accumulation in the medial temporal cortex of EMCI and LMCI, respectively, for participants with AD probability score ranges (0 ≤ AD probability score ≤ 0.05 versus 0.95 ≤ AD probability score ≤ 1.00, the ranges to which 65% of EMCI and 62% of LMCI belong ; 0 ≤ AD probability score < 0.5 versus 0.5 < AD probability score ≤ 1.00). In EMCI (**Fig. 3C**), a comparison between participants with 0 ≤ AD probability score ≤ 0.05 and those with 0.95 ≤ AD probability score ≤ 1.00 yielded a difference of 0.19 SUVR in the medial temporal lobe. In LMCI (**Fig. 3D**), the comparison of participants with low AD probability scores (0 ≤ AD probability score ≤ 0.05) and LMCI with high AD probability scores (0.95 ≤ AD probability score ≤ 1.00) yielded a difference of 0.26 SUVR in the medial temporal cortex. Whole brain voxel-wise analysis in SPM12 was performed to identify brain regions showing difference between tau deposition between MCI participants with low AD probability scores (0 ≤ AD probability score ≤ 0.05) and those with high AD probability scores (0.95 ≤ AD probability score ≤ 1.00). In EMCI (**Fig. 4A, C**) and LMCI (**Fig. 4B, D**), voxel-wise analysis identified significant group differences in the bilateral temporal lobes including the entorhinal cortex. In addition, the differences in tau deposition were more widespread in LMCI compared to EMCI (**Fig. 4**).

**Fig. 3.**
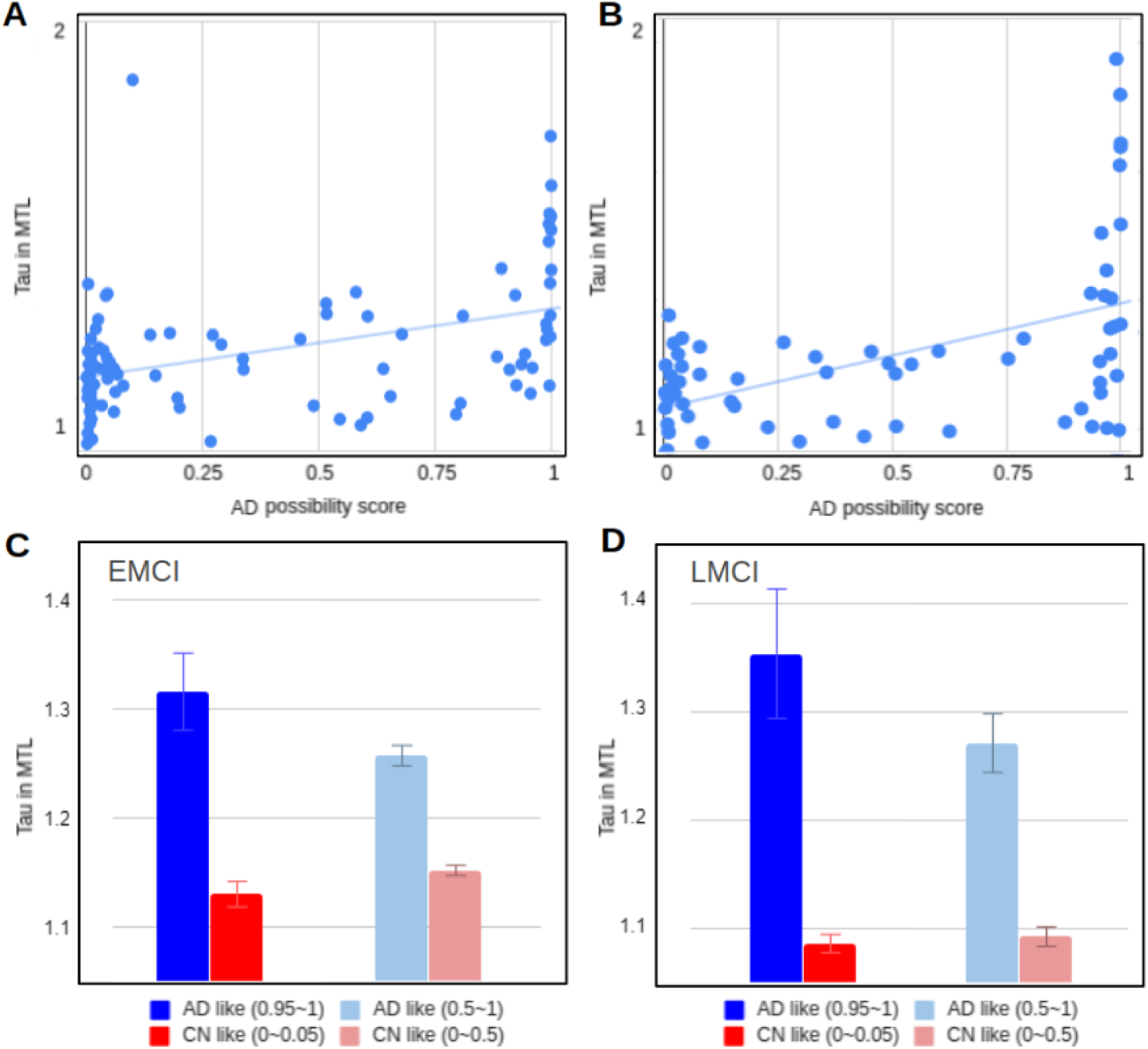
Results of scoring all the images through the classifier and comparing the scores to the tau accumulation in the MTL region. Correlation of DL score with the amount of tau accumulated in the MTL region was R = 0.43 for EMCI(A) and R = 0.49 for LMCI (B). The red and blue bar chart in C and D show the average of the tau amounts of the image with the DL score of 5% and the image of the top 95%. The charts in light red and light blue in C and D are the result of averaging the bottom 50% and top 50% of the images.

**Fig. 4.**
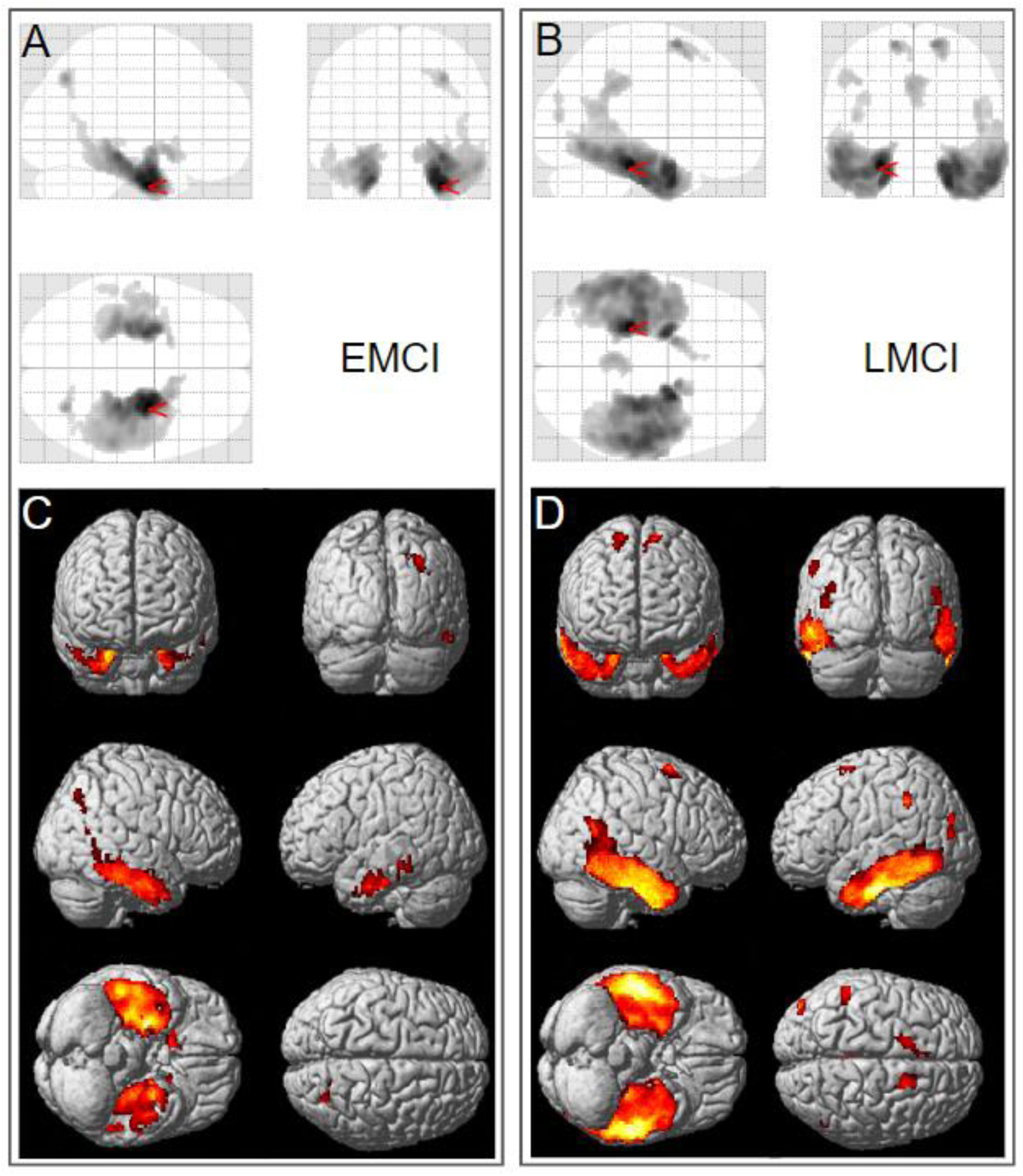
Voxel-wise difference between MCI participants with AD-like tau patterns and CN-like tau patterns defined using the 3D-CNN classifier. Significantly greater tau was observed in EMCI (A, C) and LMCI (B, D) with high AD probability (“AD-like,” 0.95 ≤ AD probability score ≤ 1.00) relative to the low AD probability group (“CN-like,” 0 ≤ AD probability score ≤ 0.05). Voxel-wise significance maps are displayed at FWE corrected p-value < 0.05; minimum cluster size (k) = 50.

## Discussion

We developed a deep learning framework for detecting informative features in tau PET for the classification of Alzheimer’s disease. After training a 3D CNN-based AD/CN classifier on 132 [^18^F]flortaucipir PET images to distinguish AD with >90% accuracy, heatmaps were generated by a LRP algorithm to show the most important regions for the classification. This model was then applied to [^18^F]flortaucipir PET images from 168 MCI to classify them into “AD similar” and “CN similar” groups for further investigation of the morphological characteristics of the tau deposition.

Maass, et al. [7] examined the key regions of *in vivo* tau pathology in ADNI using a data-driven approach and determined that the major regions contributing to a high global tau signal mainly overlapped with Braak stage III ROIs (i.e., amygdala, parahippocampal gyri and fusiform). Our deep learning-based results correspond well to the pattern reported by Maass, et al. [7] on a more limited data set.

It is noteworthy that stages III / IV can be seen in both CN and AD patients, while stages I / II are common in CN and stages V / VI are common for AD patients [3]. Thus, it is difficult to predict AD by measuring tau deposition in stage III / IV ROIs, highlighting the importance of understanding the morphological characteristics of tau. Our heatmaps, which visualized the regions driving the classification of AD and CN using deep learning on tau PET images, showed a distribution pattern similar group differences in tau deposition between AD and CN assessed using voxel-wise analysis in SPM12. This finding indicates that the deep learning classifier used the morphological characteristics of the tau distribution for classifying AD from CN. In particular, the heatmaps show that the hippocampus, parahippocampal gyrus, thalamus and fusiform gyrus were primarily used to classify AD from CN. These results support existing research showing that tau accumulation in memory-related areas play an important role in the development of AD [33, 34].

Early, accurate and efficient diagnosis of AD is important for initiation of effective treatment. Prognostic prediction of the likelihood of conversion of MCI to AD plays a significant role in therapeutic development and ultimately will be important for effective patient care. Thus, the CN vs. AD classifier was used to generate a score of whether the tau distribution in MCI participants was similar or different from that seen in AD. When the AD probability score generated by the classifier was high suggesting high similarity to AD, the MCI participants generally had the characteristic tau morphology seen in AD. In addition, we assessed applied this method to both EMCI and LMCI participants. Pearson correlation coefficients between AD probability scores and bilateral mean of SUVR in the medial temporal lobe were R = 0.43 for EMCI and R = 0.49 for LMCI. These findings indicate that the tau deposition difference between the lower 5% and upper 95% of LMCI participants was 7.1% more than the difference between the lower 5% and upper 95% of EMCI participants. Thus, the classifier determined that the tau deposition of LMCI participants is more similar to those seen in AD than that of EMCI participants. The is in line with numerous reports of biomarkers in late MCI where there is considerable overlap with early stage AD pathology.[18]

## Conclusion

Deep learning can be used to classify tau PET images from AD patients versus controls. Furthermore, this classifier can score the tau distribution by its similarity to AD when applied to scans from older individuals with MCI. A deep learning derived AD-like tau deposition pattern may be useful for early detection of disease during the prodromal or possibly even preclinical stages of AD on an individual basis. Advances in predictive modelling are needed to develop accurate precision medicine tools for AD and related neurodegenerative disorders, and further developments can be expected with inclusion of multi-modality data sets and larger samples.

## Declarations

### Ethics approval and consent to participate

Ethics approval is not required as the human data were publicly available by ADNI website, and all the data are not identifiable.

### Consent for publication

Not applicable.

### Availability of data and materials

The datasets used and analyzed during the study are available in the ADNI LONI repository, http://adni.loni.usc.edu/

## Authors’ contributions

TJ, KN, and AS: conceptualization and study design. TJ, SR: data collection. TJ: conducted the experiment and analysis. TJ, KN: drafting manuscript. TJ, KN, SR, and AS: revision of the manuscript for important scientific content and final approval.

## Acknowledgements

Data collection and sharing for this project was funded by the Alzheimer’s Disease Neuroimaging Initiative (ADNI) (National Institutes of Health Grant U01 AG024904) and DOD ADNI (Department of Defense award number W81XWH-12-2-0012). ADNI is funded by the National Institute on Aging, the National Institute of Biomedical Imaging and Bioengineering, and through generous contributions from the following: AbbVie, Alzheimer’s Association; Alzheimer’s Drug Discovery Foundation; Araclon Biotech; BioClinica, Inc.; Biogen; Bristol-Myers Squibb Company; CereSpir, Inc.; Cogstate; Eisai Inc.; Elan Pharmaceuticals, Inc.; Eli Lilly and Company; EuroImmun; F. Hoffmann-La Roche Ltd and its affiliated company Genentech, Inc.; Fujirebio; GE Healthcare; IXICO Ltd.; Janssen Alzheimer Immunotherapy Research & Development, LLC.; Johnson & Johnson Pharmaceutical Research & Development LLC.; Lumosity; Lundbeck; Merck & Co., Inc.; Meso Scale Diagnostics, LLC.; NeuroRx Research; Neurotrack Technologies; Novartis Pharmaceuticals Corporation; Pfizer Inc.; Piramal Imaging; Servier; Takeda Pharmaceutical Company; and Transition Therapeutics. The Canadian Institutes of Health Research is providing funds to support ADNI clinical sites in Canada. Private sector contributions are facilitated by the Foundation for the National Institutes of Health (www.fnih.org). The grantee organization is the Northern California Institute for Research and Education, and the study is coordinated by the Alzheimer’s Therapeutic Research Institute at the University of Southern California. ADNI data are disseminated by the Laboratory for Neuro Imaging at the University of Southern California.

The authors are grateful to Paula Bice, PhD for editorial assistance.

